# Identification of Cell-Free DNA Methylation Patterns Unique to the Human Left Ventricle as a Potential Indicator of Acute Cellular Rejection

**DOI:** 10.1101/2021.03.10.434822

**Authors:** Sabrina Pattar, Mohammad Aleinati, Fatima Iqbal, Aiswarya Madhu, Samuel Blais, Xuemei Wang, Frederic Dallaire, Yinong Wang, Debra Isaac, Nowell Fine, Steven C. Greenway

**Affiliations:** Department of Biochemistry and Molecular Biology, Cumming School of Medicine, University of Calgary, Calgary, AB, Canada; Department of Pediatrics and Alberta Children’s Hospital Research Institute, Cumming School of Medicine, University of Calgary, Calgary, AB, Canada; Department of Cardiac Sciences and Libin Cardiovascular Institute, Cumming School of Medicine, University of Calgary, Calgary, AB, Canada; Division of Pediatric Cardiology, Department of Pediatrics, Faculty of Medicine and Health Sciences, Université de Sherbrooke and Centre de Recherche du Centre Hospitalier Universitaire de Sherbrooke, Sherbrooke, QC, Canada; Alberta Precision Laboratories and Department of Pathology & Laboratory Medicine, Cumming School of Medicine, University of Calgary, Calgary, AB, Canada

**Keywords:** epigenetics, DNA methylation, heart, transplantation, cell-free DNA, rejection

## Abstract

Increased levels of donor-derived cell-free DNA (dd-cfDNA) in recipient plasma have been associated with rejection after transplantation. DNA sequence differences have been used to distinguish between donor and recipient but epigenetic differences could also potentially identify dd-cfDNA. This pilot study aimed to identify ventricle-specific differentially methylated regions of DNA (DMRs) that could be detected in cfDNA. We identified 24 ventricle-specific DMRs and chose two for further study, one on chromosome 9 and one on chromosome 12. The specificity of both DMRs for the left ventricle was confirmed using genomic DNA from multiple human tissues. Serial matched samples of myocardium (n=33) and plasma (n=24) were collected from stable adult heart transplant recipients undergoing routine endomyocardial biopsy for rejection surveillance. Plasma DMR levels increased with biopsy-proven rejection grade for individual patients. Mean cellular apoptosis in biopsy samples increased significantly with rejection severity (2.4%, 4.4% and 10.0% for ACR 0R, 1R and 2R, respectively) but did not show a consistent relationship with DMR levels. We identified multiple DNA methylation patterns unique to the human ventricle and conclude that epigenetic differences in cfDNA populations represent a promising alternative strategy for the non-invasive detection of rejection.

## Introduction

All cells undergoing programmed cell death (apoptosis) release fragments of double-stranded genomic DNA (gDNA) into the circulation. Plasma cell-free DNA (cfDNA) has shown promise as a minimally-invasive biomarker for the detection of rejection after solid organ transplantation although many questions regarding the production and clearance of cfDNA remain unanswered.^1–3^ To date, the identification of donor-derived cfDNA has been based on the quantification of polymorphic single nucleotide polymorphisms (SNPs) that differ between the donor and recipient with or without donor genotyping.^4–6^ While the requirement for *a priori* donor genotyping has limitations associated with cost and feasibility, especially in situations where the donor is deceased, alternative methods are subject to assumptions and estimations of genotype frequencies that may not accurately represent the donor.^4–6^ As such, epigenetic modifications (e.g. DNA methylation) that remain intact in cfDNA represent a potential alternative methodology to identify specific populations of cfDNA within the complex mixture found in the plasma.^7, 8^ Unique tissue-specific DNA methylation patterns known as differentially methylated regions (DMRs) have been identified and leveraged as markers of organ injury^9^ but are understudied in cardiovascular disease^10^ and remain unexplored in transplantation.

In this pilot study, we sought to develop a pipeline to identify novel alternative markers of myocardial injury based on DNA methylation. From 24 putative candidates, we studied two ventricle-specific DMRs detectable in plasma cfDNA. Using matched samples of plasma and myocardium from adult heart transplant recipients undergoing routine endomyocardial biopsy (EMB), we then evaluated the links between levels of ventricle-specific cfDNA, grade of acute cellular rejection (ACR) and myocardial apoptosis.

## Material and Methods

### Patient data

This study was approved by the Conjoint Health Research Ethics Board at the University of Calgary. All participants provided written consent. Adult heart transplant recipients followed at a single centre and undergoing a scheduled EMB were recruited for study participation. All patients were at least 3 weeks post-transplant, had no evidence of cardiac dysfunction, antibody-mediated rejection (AMR) or cardiac allograft vasculopathy (CAV) and had at least three serial biopsies with tissue available and matching plasma samples. The right ventricular (RV) myocardial tissue was obtained from paraffin-embedded blocks that had been used for pathology grading by a single pathologist according to the current ISHLT guidelines.^11^ Biopsies graded as having ACR 2R were independently reviewed by a second pathologist to confirm the diagnosis as per institutional policy.

### Isolation of cfDNA

Blood (8-10 mL) was collected in Streck BCT tubes immediately prior to the EMB. Within 4 hours of collection, whole blood was centrifuged at 1900 x g at room temperature for 10 minutes. The plasma fraction was then centrifuged at 13,000 RPM for 16 minutes at 4°C. The resulting supernatant was then stored at −80°C. Isolation of cfDNA from 2 mL of plasma was performed using the semi-automated MagNA Pure 24 System (Roche) according to the manufacturer’s instructions.

### *In silico* identification of candidate ventricle-specific DMRs

Public methylomes for various human tissues and cells were obtained from the NIH Roadmap Epigenomics^12^ and the Blueprint Epigenome^13^ projects. Sorted bedgraph input files were created for the software package Metilene^14^ which performed comparisons between the methylomes of the ventricular tissue (combined right (n=1) and left (n=2) ventricles), pooled non-ventricular tissues (n=19) and pooled hematopoietic cells (n=14). Metilene identified regions that were significantly differentially methylated (mean difference >10%) based on a Mann-Whitney U test (alpha = 0.05). Additional parameters for each DMR included a minimum of 3 CpG sites, <25 base pairs (bp) separating each of the CpG sites and a total length of <100 bp. A second filtering step identified DMRs with a mean methylation difference >50% for non-ventricular tissues and >80% for hematopoietic cells. Comparisons between groups (ventricles vs. hematopoietic cells or non-ventricular tissues) were made separately to enable stringent exclusion of DMRs in peripheral lymphocytes which account for the majority of cfDNA in the circulation. The filtering cutoffs chosen resulted in a sufficient number of DMRs within each interrogated group to enable the identification of DMRs that were common to both groups. Shared DMRs on autosomal chromosomes (sex chromosomes were excluded to avoid dosage effects) identified from the comparison of ventricular and non-ventricular tissue methylomes (n=255) and from ventricular tissues compared to hematopoietic cell methylomes (n=2,060) provided a total of 24 candidate DMRs (Figure 1).

**Figure 1.**
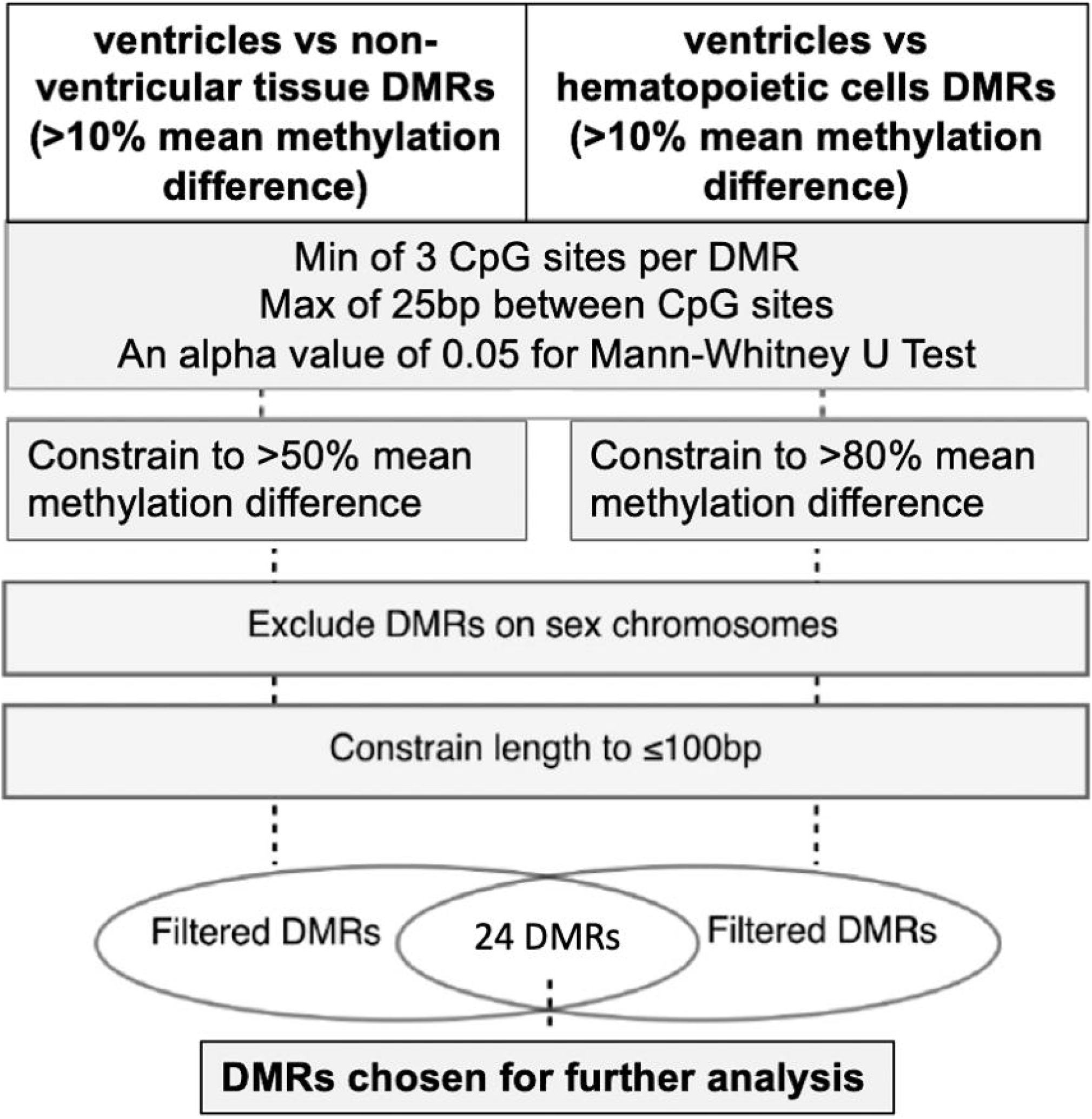
DMR identification flow chart. Publicly available methylomes were compared using Metilene to identify DMRs that were unique to the ventricles.

### Primer design

For 10 candidiate DMRs with the highest ventricle-specificity, sequences containing the DMR of interest and 100 bp upstream and downstream were taken from a methylome of the left ventricle (GSM1010978) generated by the NIH Roadmap Epigenomics project. These sequences were used as input for MethPrimer^15^ to generate primers for amplification of the DMRs from bisulfite-converted gDNA. Primers did not contain CpG islands and contained at least two non-CpG cytosines. Total product size was constrained to <150 bp to enable amplification from cfDNA. From these, two DMRs, on chromosomes (Chr) 9 and 12, were chosen for further study as these regions amplified readily with no off-target products.

### PCR amplification of bisulfite-converted DNA

A commercial panel (Zyagen) of gDNA from human brain, colon, esophagus, small intestine, kidney, liver, lung, pancreas, stomach, skin, spinal cord, skeletal muscle, spleen and thymus was used to compare methylation levels for the Chr 9 and Chr 12 DMRs across tissues in comparison to left ventricle. Patient cfDNA and panel gDNA was bisulfite converted using the Epitect Bisulfite Kit (Qiagen) with elution performed twice using 20 μL of buffer EB warmed to 56°C to improve yields. Bisulfite-converted DNA was used for PCR amplification of the two DMRs. Products were visualized on a 3% TAE agarose gel and quantified by TapeStation (Agilent).

### Next-generation sequencing

For quantification of methylation, PCR amplicons were pooled in equimolar concentrations for sequencing on an Illumina MiSeq. Libraries were sequenced using the MiSeq reagent kit v3 (150 cycles) to produce 2 × 75 bp paired-end reads. The quality of each sequenced pool was assessed using FastQC.^16^ The bisulfite sequencing plugin for the CLC Genomics Workbench (Qiagen) was utilized to analyze the reads which were mapped to the hg19 reference genome with a minimum acceptable alignment of >80%. Methylation levels were determined for each individual CpG site within each DMR and mean methylation levels were calculated. For DMR analysis, only those patients with available matched samples (myocardium and plasma) were included.

### Determination of cfDNA concentrations

The concentration of cfDNA (ng/mL) extracted from 2 mL of patient plasma was divided by 0.00303 ng (mass of a single human genome) to determine the total concentration of cfDNA (copies/mL) in recipient plasma.^9^ To determine the absolute concentration of each DMR, the copies/mL was multiplied by the fraction of unmethylated molecules within the given pool of amplicons as determined from the analysis of the bisulfite-sequenced reads.

### Analysis of myocardial apoptosis

Terminal deoxynucleotidyl transferase dUTP nick end labelling (TUNEL) staining (Promega) was used to visualize fragmented gDNA within paraffin-embedded RV tissue sections. Slides were prepared and TUNEL staining performed as per the manufacturer’s instructions. Tissue slides were also stained with DAPI (4′,6-diamidino-2-phenylindole, Invitrogen) to visualize the total number of nucleated myocardial cells within each section. Stained tissue slides were imaged at 10X magnification using a spinning disk confocal super resolution microscope (Olympus). A green fluorescent signal was observed at 488 nm in regions that contained apoptotic cells while DAPI-stained nuclei emitted a blue signal at 405 nm. Images were analyzed using ImageJ and the Fiji plugin. Analysis of the tissue sections was performed while blinded to the results of the pathology ACR scoring.

### Statistical analysis

To evaluate the association between rejection grade and cellular apoptosis and between rejection grade and copies/mL of ventricle-specific cfDNA, we used multilevel linear regression. This allowed adjustment for the potential cluster effect of comparing multiple samples per biopsy and repeated biopsies for each subject. We used an iterative approach to test several models and evaluated the necessity of including random effects for each level. The final model consisted of a three-level hierarchical linear regression of log-transformed apoptosis percentages according to rejection grade. Cellular apoptosis in the tissue sections showed a log-normal distribution and, therefore, the apoptosis data are presented as geometric means and standard deviations. For the cfDNA data, the mean of the 0R measurements were used to compare the effect of rejection on the log increase of cfDNA concentration compared to baseline. Statistical analyses were performed using SAS software version 9.4 (SAS Institute).

## Results

### Ventricle-specific DMRs identified *in silico*

Methylomes obtained from the NIH Roadmap Epigenomics and the Blueprint Epigenome projects included hematopoietic cells (n=14), various extra-cardiac tissues (n=19) and the left (n=2) and right ventricles (n=1) of the heart. Since our analysis of the 3 available ventricular samples indicated that the human left and right ventricles were not significantly epigenetically distinct, these datasets were combined for this analysis. By comparing the ventricular methylomes to the 19 tissue methylomes, 255 DMRs were identified. The majority (95.8%) of these were hypomethylated in the ventricles relative to the other tissues. The comparison of the ventricular methylomes to the 14 methylomes of hematopoietic cells identified 2,060 DMRs that were either hypomethylated (37.2%) or hypermethylated (63.8%) in the ventricles. A total of 24 DMRs were common to both. From these 24 putatively ventricle-specific DMRs, two displayed the greatest ventricular specificity and amplified specifically and were therefore chosen for further study. The first DMR was identified on Chr 9 (position Chr 9:101,595,534-101,595,574) and contained 3 CpG sites over a span of 40 bp. The second DMR contained 6 CpG sites within 44 bp and was found on Chr 12 (position Chr 12:106,132,052-106,132,096). Interestingly, although we found that the methylation patterns in these regions are unique to ventricular cells relative to other tissues and cells, these DMRs are not located within or adjacent to genes directly associated with cardiac function or development. Based on the hg19 reference genome, the DMR on Chr 9 was found to be located within the gene body of *GALNT12*, a protein-coding gene associated with colorectal cancer, and the Chr 12 DMR was found to be located within the gene body of *CASC18*, which is a non-protein coding gene that has been identified as a potential cancer predisposition locus.

### *In vitro* confirmation of ventricular specificity

To confirm our *in silico* findings, we interrogated gDNA from left ventricle (LV) and human tissues encompassing all three developmental germ layers (n=14). After amplification from bisulfite-converted LV gDNA, the three individual CpG sites within the Chr 9 DMR were found to be methylated in 18-26% of the sequenced amplicons with an average methylation level of 22.7% and the six CpG sites within the DMR on Chr 12 were found to the methylated in 16-22% of the sequenced amplicons with an average methylation level of 21%. These levels of methylation were lower than those predicted from our *in silico* analysis. As shown in Figure 2, in comparison to other tissues, both DMRs were most unmethylated in the LV (77.3 ± 0.04% for Chr 9 and 80.0 ± 0.02% for Chr 12) although pancreas and lung were also mostly unmethylated at the Chr 9 locus.

**Figure 2.**
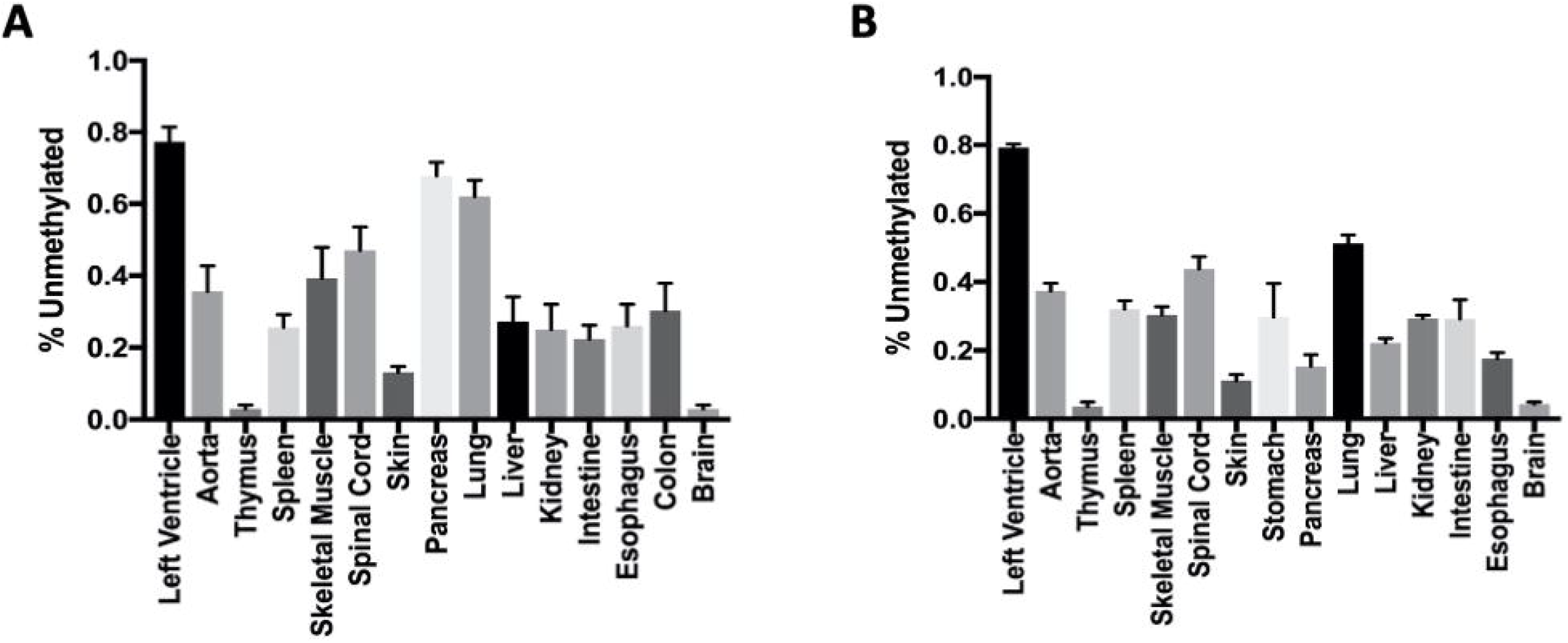
Methylation levels (% unmethylated) for bisulfite-converted gDNA isolated from various human tissues as determined *in vitro*. (A) Tissue methylation levels for Chr 9 DMR. (B) Tissue methylation levels for Chr 12 DMR. Data shown are average methylation levels ± SD.

### Performance of ventricle-specific DMRs in detecting significant rejection

To assess the ability of our two DMRs to detect significant ACR, serial plasma samples obtained at the time of routine EMB were tested. Both DMRs showed variable levels that increased with the rejection grade independently assigned by the pathologist (Figure 3). Highest levels of both DMRs were seen with the significant 2R events and lowest levels were present with no (0R) or mild (1R) ACR. The concentration of both DMRs remained relatively stable across the 0R samples, ranging from 1,100-2,987 copies/mL. In contrast, samples collected at the time of a 1R ACR event showed the most variability, ranging in concentration from levels below those seen in 0R (370-660 copies/mL) to levels approaching those seen in grade 2R ACR (11,480-17,710 copies/mL). Interestingly, we noted an increase in the concentration of both DMRs within the plasma of patient 3 prior to a moderate rejection event (Figures 3B and 3E). DMR concentrations also appear to be patient-specific with differences between individuals at baseline (0R) and also for the 1R and 2R events. Overall, levels of cfDNA during ACR 2R events were significantly increased (p = 0.03) in comparison to baseline 0R levels for the Chr 12 DMR (Figure 3F) but were not significant for the Chr 9 DMR (p = 0.077) (Figure 3C). In general, total cfDNA levels in recipient plasma correlated poorly with rejection severity.

**Figure 3.**
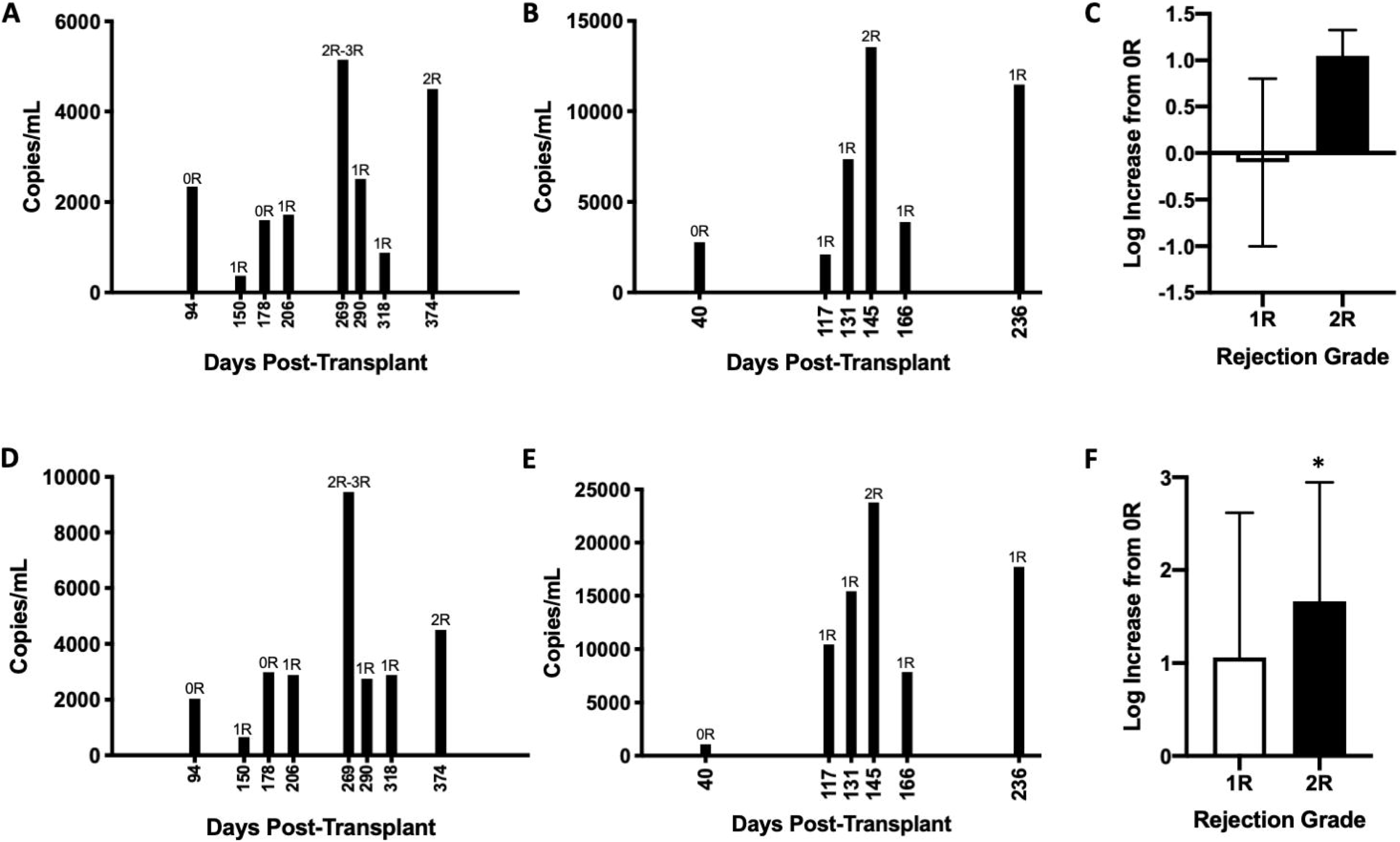
Serial plasma DMR concentrations and rejection grades. (A) Levels of plasma Chr 9 DMR for patient 1 at EMBs taken during days 94-374 post-transplantation. (B) Levels of plasma Chr 9 DMR for patient 3 from days 40-236 post-transplantation. (C) Log increase in copies/mL for the Chr 9 DMR for the 1R and 2R rejection grades compared to the mean level of the 0R samples. There were no significant differences in the plasma concentration of cfDNA associated with either the 1R (p = 0.69) or 2R (p = 0.077) rejection grades for the Chr 9 DMR. (D) Levels of plasma Chr 12 DMR for patient 1 from days 94-374 post-transplantation. (E) Levels of plasma Chr 12 DMR for patient 3 from days 40-236 post-transplantation. The associated pathology rejection grade (ACR 0R, 1R or 2R) is indicated for each sample. (F) Log increase in copies/mL for the Chr 12 DMR for the 1R and 2R rejection grades compared to the mean level of the 0R samples. There was a significant (p = 0.03) increase in the plasma concentration of cfDNA associated with a 2R rejection grade but not for the 1R rejection grade (p = 0.2).

### Myocardial apoptosis is increased in acute cellular rejection

Serial EMB samples obtained from six individual adult heart transplant patients were used for fluorescence microscopy. A total of 33 separate biopsies (n=8 for ACR 0R; n=17 for ACR 1R; n=8 for ACR 2R) were included in the analysis. DAPI staining determined the total number of nuclei within the biopsied tissue section and the number of cells undergoing apoptosis was indicated by the presence of fragmented DNA as determined by TUNEL staining. Cells that displayed colocalization of DAPI and TUNEL were considered to be apoptotic.

Rejection grade assigned by the pathologist was associated with the number of areas within the biopsied tissue that displayed elevated levels of apoptosis (Figure 4A-D). In general, those samples without significant ACR (ISHLT grade 0R) had the fewest number of apoptotic cells (<12%) with no focal areas of cell death. In contrast, samples that were assigned a rejection grade of 1R (single area of lymphocyte infiltration) displayed a single region of elevated apoptosis representing 20-35% of the total cells and a variable degree of background apoptosis in the remaining tissue. Samples assigned a rejection grade of 2R (two areas of lymphocyte infiltration) all displayed two areas in which 20-34.3% of the myocardial cells co-stained for both TUNEL and DAPI. The adjusted mean percentage of myocardial cells that stained positively for TUNEL was observed to be 2.4% (CI 1.6-3.6), 4.4% (CI 2.8-6.9) and 10.0% (CI 5.8-17.2) for rejection grades 0R, 1R, and 2R, respectively (Figure 4E). Thus, the mean percentage of apoptotic cells increased as the severity of rejection increased and differences between the groups were statistically significant (0R vs. 1R, p=0.01; 1R vs. 2R, p=0.0004 and 0R vs. 2R, p<0.0001). Patient 2 (Figure 4B) had several biopsies that were assigned a rejection grade of 1R and in each a single area of elevated apoptosis ≥10% (ranging from 11.8-17.6%) was noted. In this patient, the degree of tissue apoptosis was generally similar to that seen in the 0R samples but distinguished by a single region of increased cell death. However, in patient 3 (Figure 4C), the 1R samples were more variable.

**Figure 4.**
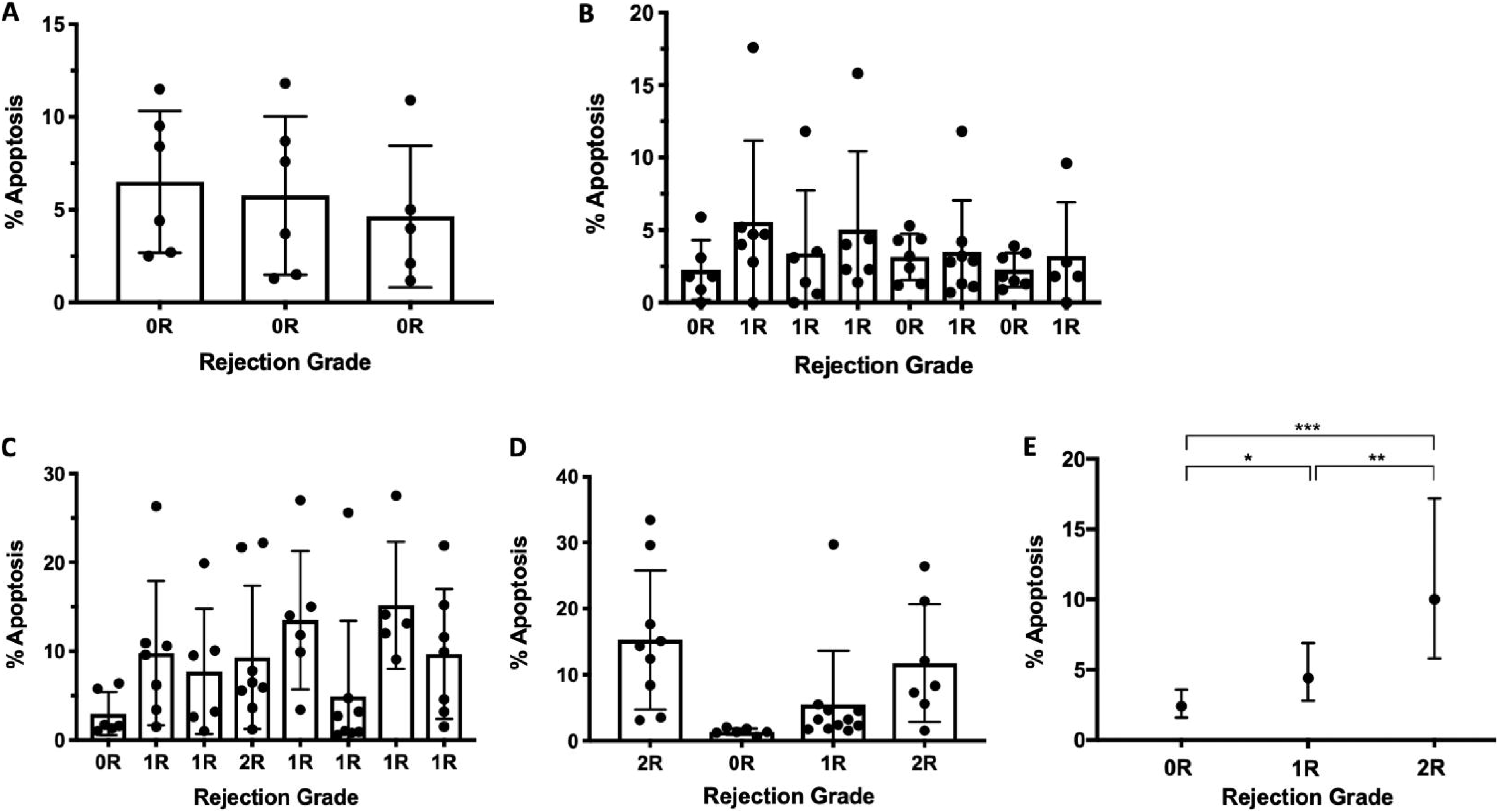
Percentage of cells undergoing apoptosis as defined by number of cells co-localizing DAPI and TUNEL staining. (A) Patient 1 with no significant rejection episodes. (B) Patient 3 with no serious rejection episodes. (C) Patient 4 with multiple episodes of mild rejection and one significant (2R) rejection event. (D) Patient 2 with two episodes of significant rejection. Data are mean and SD with individual data points shown as circles representing sections analyzed for each separate biopsy sample. (E) Data summary with mean percentage of TUNEL-positive myocardial cells at each rejection grade. The adjusted mean proportion of myocardial cells that stained positively for TUNEL from 33 separate biopsies (n=8 for ACR 0R, n=17 for ACR 1R and n=8 for ACR 2R) from 6 individual patients. *p=0.01, **p=0.004, ***p<0.0001. Data are means with bars representing 95% confidence intervals.

### Relationship between ventricle-specific DMRs and myocardial apoptosis

We attempted to associate levels of ventricle-specific cfDNA with mean levels of tissue apoptosis in serial EMB samples from three individual patients with available data (Figure 5). The relationship between apoptosis and levels of the two DMRs were variable for each patient, showing either a negative or positive fit, which may be an effect of the small sample size and/or the large variation in the concentration of the DMRs in samples assigned a rejection grade of 1R. Notably, the two DMRs behaved similarly in each patient.

**Figure 5.**
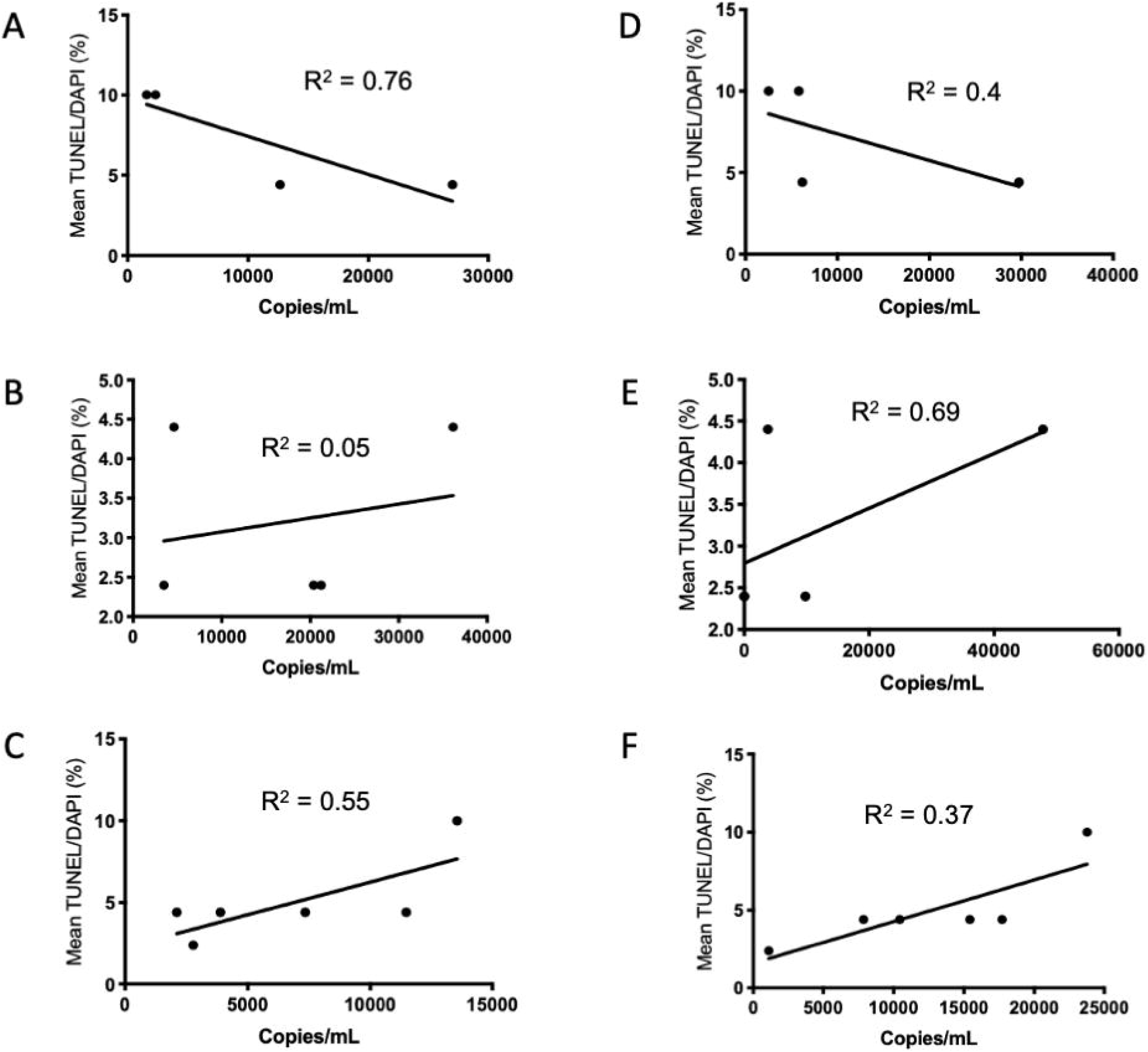
Association of DMR concentration (copies/mL) with cellular apoptosis. (A-C) Relationship between myocardial apoptosis and the Chr 9 DMR for patients 2-4. (D-F) Relationship between myocardial apoptosis and the Chr 12 DMR for patients 2-4.

## Discussion

DNA methylation plays a critical role in dictating gene expression and controlling cellular differentiation and specialization.^17, 18^ The observations that DNA methylation patterns could identify tissue-specific cfDNA populations^7–9^ led us to identify ventricle-specific patterns using an unbiased approach and publicly-available methylomes whereby entire methylomes were interrogated instead of selecting regions previously established as being associated with the patterning and development of the myocardium. This approach allowed for the identification of unique or highly specific biomarkers that may have been otherwise undetected due to a lack of genome annotation. However, examining regulatory regions of unique myocardial genes represents an alternative approach to the discovery of ventricle-specific DMRs. Methylomes from hematopoietic cells were also assessed as the natural turnover of these cells is the primary contributor to the total plasma cfDNA pool and therefore contributes most to background noise when measuring ventricle-derived cfDNA.^19^

From a total of 24 DMRs, candidates on Chr 9 and 12 were selected for further analysis. *In vitro* testing of these two DMRs revealed less than expected specificity for the ventricle. This may be related to the small number of available methylomes used to identify these DMRs but also suggests the importance of confirming all *in silico* predictions. There was reasonable agreement between the concentration of the ventricle-specific biomarkers and the severity of rejection with the most variability noted within the 1R samples. This may be related to the combination of the old grades of 1A, 1B and 2 into a single category.^11^ The heterogeneity of the 1R specimens was also present in our apoptosis data. DMR concentrations also appear to be patient-specific with relatively large differences (>1,000 copies/mL) between individuals at baseline (0R) and also for the 1R and 2R events. This suggests that DMRs are not generic but may be useful as personalized indicators that can be followed serially (as represented in Figure 3) with increases suggesting the implementation of additional testing (e.g. an EMB) or changes in management. As such, the use of these DMRs may be the most advantageous for those who have displayed consistent biopsy results that indicate stable quiescence, whereby the adverse effects of the EMB outweigh the benefits of performing the procedure.

ACR results in the death of myocardial cells through the accumulation of CD4+ and CD8+ T-cells in the interstitial space of the allograft which initiates apoptosis either through the release of cytotoxic granules or through direct binding to FasR on the target cell.^20^ The presence of increased cell death has been documented in both cardiomyocytes^21–23^ as well as in inflammatory and interstitial cells.^24, 25^ With regards to myocardial apoptosis in our study, the number of areas of lymphocyte infiltration associated with each rejection grade were consistent with the number of areas that demonstrated elevated apoptosis within the ventricular tissue sections. The amount of apoptosis occurring outside of focal areas of lymphocyte infiltration was especially variable within individual ACR 1R samples. Given that total cfDNA in the plasma arises from all cells and tissues in the body and can be affected by multiple physiological (e.g. strenuous exercise, fever)^26, 27^ and non-physiological stressors (e.g. surgery)^28, 29^ it is perhaps not surprising that it was insufficient to detect organ-specific injury.

Our study has several important limitations. As a pilot study assessing feasibility, we studied relatively few patients although we did have multiple and matched serial plasma and tissue samples for each patient. Replication of our findings in a larger cohort of patients, especially during severe (2R or 3R) ACR events, will be important to validate the generalizability of our findings. Alterations in our filtering criteria for the selection of ventricle-specific DMRs are also possible and may be important when identifying tissue-specific DMRs in other organs. We also did not identify the specific cell type(s) undergoing apoptosis in our biopsy samples. The literature suggests that both inflammatory cells and cardiomyocytes undergo apoptosis during rejection events, with several reports describing that it is mostly inflammatory cells that are undergoing apoptosis during rejection.^24, 25^ Our observation that the number of clusters of apoptotic cells are equivalent to the number of clusters of lymphocytes are therefore consistent with these previous observations. If lymphocytes are the cells predominantly undergoing apoptosis during rejection this may explain the poor correlation between the degree of apoptosis and plasma levels of our DMRs. We did not specifically evaluate the methylation levels of our DMRs in B cells or inflammatory cells but this could be done in the future. Finally, the TUNEL assay that we used to assess cell death is non-specific and labels all fragmented DNA and includes those cells undergoing either apoptosis or necrosis.^30, 31^ However, both apoptosis^25^ and necroptosis^32, 33^ are involved in acute and chronic cardiac allograft rejection. In the future, it will be useful to evaluate phosphorylated mixed lineage kinase domain-like protein (pMLKL) in our tissue sections to assess the contribution of necroptosis to cell death along with a more specific marker for apoptosis (e.g. cleaved caspase-3).^30, 34^ It will also be useful to evaluate our DMRs in other settings of allograft injury (e.g. AMR, CAV) or myocardial injury resulting from ischemia or heart failure.^10^

## Conclusions

In summary, we introduce the possibility of using ventricle-specific DMRs as a minimally-invasive cfDNA-based assay for the detection of ACR following heart transplantation in adults and as a potential alternative to SNP-based analyses. The pipeline we have established for the identification of ventricle-specific DMRs can be easily modified and applied to any solid organ and may even have relevance in hematopoietic stem cell transplantation. We also demonstrate that EMB samples from patients labelled 1R according to the current ISHLT grading system are highly heterogeneous based on both tissue apoptosis as well as the release of cfDNA derived from the transplanted ventricles. The heterogeneity of the 1R samples suggests an opportunity for future research to distinguish subtypes of grade 1R ACR that may merit different management strategies.

## Acknowledgements

We would like to thank Leo Dimnik in the Medical Genetics Laboratory at the Alberta Children’s Hospital for assistance in the automated isolation of plasma cfDNA and the nurses and physicians of the adult heart transplant program at the Foothills Hospital for assistance in blood collection.

## Author Contributions

SP, MA, FI, AM and XW performed experiments and acquired data. SB and FD performed the statistical analysis of the tissue apoptosis data. SP, MA and FI performed all other data analysis. SP, FI, DI and NF recruited patients and collected plasma samples. YW provided the pathology samples. SP, MA and SCG conceived and designed the experiments. SP and SCG wrote the manuscript. SCG provided funding. All authors read and approved the manuscript.

